# New Zealand spider orchid (*Corybas trilobus*) exploits the reproductive behaviour of fungus gnats (Diptera: *Mycetophila*) for pollination

**DOI:** 10.64898/2026.10.26.762021

**Authors:** CA Lehnebach, AJ Zeller, R Keyzers, J Kasper, P Garnock-Jones

**Affiliations:** Museum of New Zealand Te Papa Tongarewa. PO BOX 467. Wellington, New Zealand; Callaghan Innovation. PO BOX 31310. Lower Hutt, New Zealand; School of Chemical and Physical Sciences, and Centre for Biodiversity and Restoration Ecology, Victoria University of Wellington, PO BOX 600. Wellington, New Zealand; School of Biological Sciences, Victoria University of Wellington, PO BOX 600. Wellington, New Zealand

**Keywords:** brood-site deception, *Corybas*, Diptera, fungus gnat, *Mycetophila*, mimicry, mushroom, New Zealand, oviposition, pollination, *rendezvous-*site attraction, Orchidaceae

## Abstract

Due to a reduce pollinator diversity, plant-pollinator interactions in remote habitats are commonly unspecialised. In New Zealand (NZ), many plants exhibit a generalist pollination syndrome where flowers are open-access and pollinated by a range of insects. Orchids in the genus *Corybas* seem to be the exception. It is believed they mimic mushrooms and their pollination occur by brood-site deception. This study aims to uncover whether this pollination strategy operates in *Corybas* Remutaka.

We used hand-pollination experiments to determine the breeding system of this orchid. SEM observations and neutral red staining were used to study the floral micro-morphology and location of scent glands, respectively. Pollinators’ behaviour was observed, and filmed, in the field and *ex situ*. Olfactory signals were assessed using solid-phase micro-extraction and solvent extractions, and visual cues by photography with a UV sensor.

Only hand self- and cross-pollinated flowers formed capsules, while natural fruit-set was low. Male and female individuals of the endemic fungus gnat *Mycetophila phyllura* (Diptera) were the only pollinators observed. Both sexes can remove and deposit pollinia. None of the typical fungal compounds, or other compounds linked to brood-site deception, were detected in the orchid but a prevalence of alkane, sesquiterpene and fatty aldehydes. Some of these compounds were also detected in a mushroom used by fungus gnats for mating and ovipositioning. Features in the labellum surface and base (e.g. exit holes) and UV reflectance may help directing movement of the gnats within the flower.

*Corybas* Remutaka is a self-compatible, nectarless, pollinator-dependent orchid that relies on a mix of volatiles and micro-morphological features to attract a specific pollinator. The orchid exploits the reproductive instinct of male and female fungus gnats to secure its pollination. This deceptive pollination strategy combines features of *rendezvous* site pollination and brood-site deception.

## INTRODUCTION

Specialised pollination systems and species-specific interactions are unusual in isolated environments (Olesen and Jordano 2002). For instance, the occurrence of self-pollination and self-compatibility is a common feature across the flora of oceanic islands (Bernardello *et al*., 2001; Newstrom and Robertson, 2005; Micheneau *et al*., 2008) with some species exhibiting a more generalised pollination system than their mainland conspecific (Micheneau *et al*., 2008; Martén-Rodríguez *et al*., 2015). The relaxation of plant pollinator interactions in remote habitats has normally been attributed to reduce pollinator diversity (Carlquist, 1974; Inouye, 1993) and the over-representation of unpredictable, imprecise and inefficient pollinators such as Coleoptera and Diptera (Proctor *et al*., 1996; Faegri and van der Pijl, 1979). In the New Zealand (NZ) Archipelago, adaptation to such unusual pollinating fauna has resulted in the abundance of plants with small, open-access white or pale flowers that are dish or bowl shaped (Newstrom and Robertson, 2005). These floral traits are common in plants with a generalist pollination syndrome where flowers are visited by a range of insect pollinators (Newstrom and Robertson, 2005). However, Newstrom and Robertson (2005) caution this trend must not be overstated and they predict specialisation may occur in other insect–pollinated plants with closed-access blossoms, such as orchids.

With more than 110 species, Orchidaceae are one of the top 10 largest families of flowering plants in NZ. They are found across the entire country, including many small offshore islands and sub-Antarctic islands, occupying diverse habitats from coastal to forest and alpine settings. Most are terrestrial; with only nine species grow as epiphytes on trees, or rarely as lithophytes. Despite orchids being a conspicuous element of the NZ flora, little is known about their biology or interactions with other organism such as pollinators or mycorrhizal fungi. Although there is a plethora of photographic records of orchid-insect interactions published in the NZ native orchid journal (http://www.nativeorchids.co.nz/Journals/Journals.htm), few studies have investigated in detail their pollination strategies and breeding systems. To date, unspecialised pollination by generalist insects has been documented in four epiphytic species (Lehnebach and Robertson, 2004), self-pollination and delayed self-pollination reported in two terrestrial species (Lehnebach *et al*., 2005), and pollination by sexual deception suggested, but not demonstrated, in six species of greenhood orchids (*Pterostylis*) (Lehnebach *et al*., 2005; Bodley *et al*., 2016; Thalwitzer *et al*., 2018). Finally, brood-site deception has been suggested to occur in *Corybas* (Dafni and Bernhardt, 1990) but never fully confirmed.

In a brood-site deception system, gravid females are lured by olfactory signals to flowers that mimic oviposition sites (Urru *et al*., 2011). Flies and beetles are the main victims of this deceptive strategy and the smell produced by these flowers, which is also detectable by the human nose, resembles that of oviposition substrates such as dung and urine, carrion and rotten flesh, fermenting fruits and yeast, and fungi (Urru *et al*., 2011). Claims of brood-site deception in *Corybas* are likely based on historical observations. For instance, Thomson (1927) stated that NZ *Corybas* are nectarless, incapable of self-pollination and that they are likely pollinated by small Diptera. Fuller (1994) observed multiple female fungus gnats from different species each visiting a specific *Corybas* species. Most recently, studies of the NZ endemic *Corybas cheesemanii* found no compelling evidence of brood-site deception in that particular species (Kelly *et al*., 2013; Kelly and Gaskett 2014).

*Corybas* includes more than 150 species distributed across the Himalayas to southern China, the Malay Archipelago, Polynesia, New Guinea and Australia (Jones *et al*., 2002; Lyon, 2014) and it is one of the most common and speciose orchid genera in NZ (Schönberger *et al*., 2019). Currently, there are 22 species listed for NZ (Schönberger *et al*.; 2019) and at least four taxonomically undetermined entities (de Lange *et al*.; 2018) that are the focus of current taxonomic research by the first author. Some of these potentially new species are already considered of conservation concern, as populations are small or restricted to a single site. In NZ, and with a few exceptions, these orchids are commonly known as spider orchids. This is likely due to their long and filiform sepals and petals normally pointing upwards and sticking well above the leaf litter (Fig. 1A). One of the most widespread and morphologically variable species of spider orchid in NZ is *Corybas trilobus*. Lehnebach *et al*. (2016) segregated five species from within *C. trilobus* and stated the taxonomic status of six other entities, currently known by informal tag names only, needed further research. One of these entities is *Corybas* Remutaka, a late-winter flowering orchid endemic to NZ and restricted to southern beech (Nothofagaceae) forest in the southern North Island and the northern South Island. Similar to other *Corybas* species in NZ (Upson, 2014) and Australia (e.g. Kuiter and Findlater-Smith, 2017), micro-dipterans commonly known as fungus gnats, and insect eggs, have been found inside its flowers. Just as in *C. cheesemanii*, these observations have supported claims that pollination of this orchid occurs by brood-site deception.

**Figure 1:**
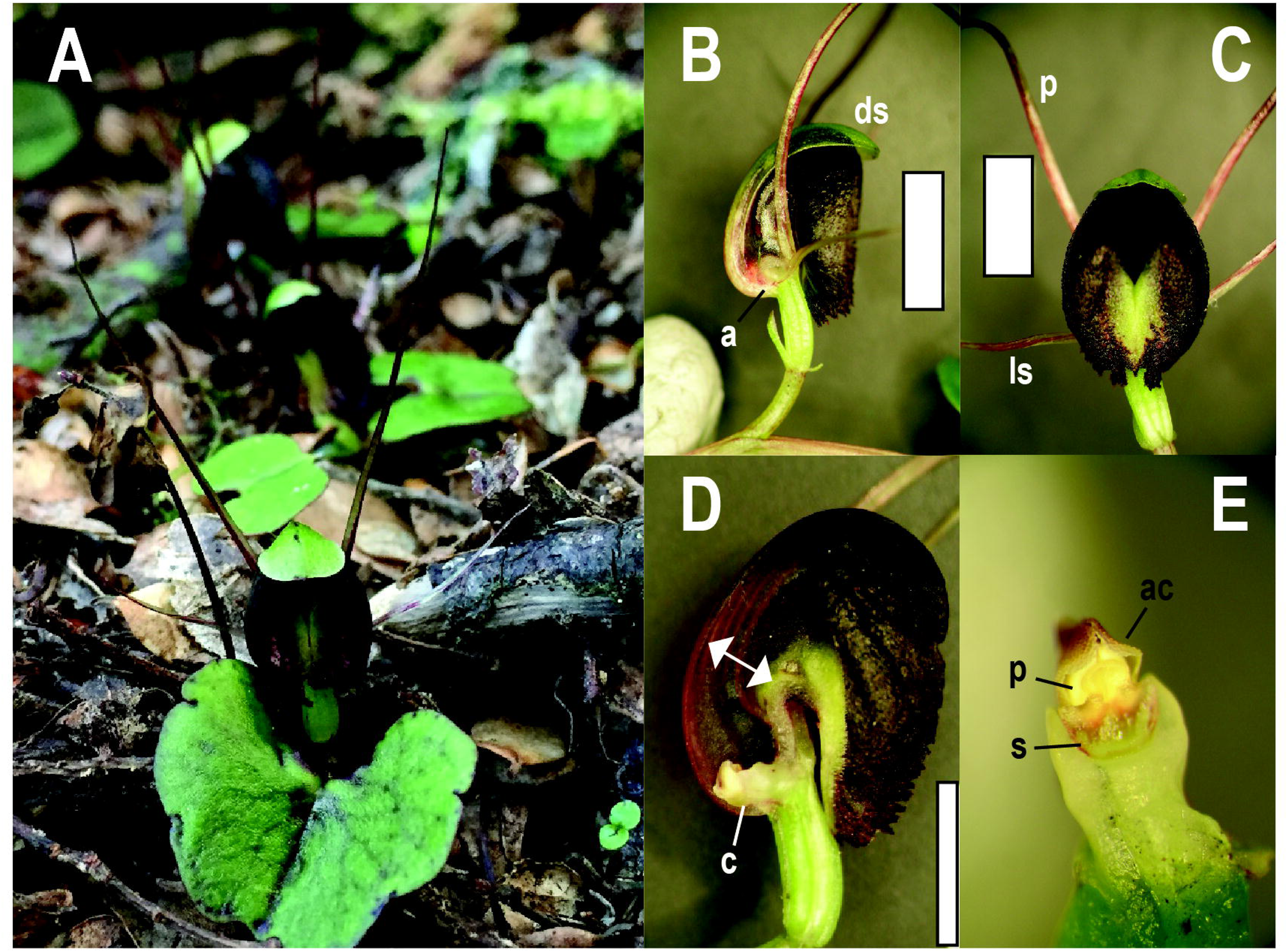
*Corybas* Remutaka and details of its flower. A: Plant in the wild. B: Side view of the flower with auricle (a) and dorsal sepal (ds). C: Frontal view of the flower with lateral sepals and petals (ls and p). D: Internal view of the flower with column (c) at the base. Arrows points to the walls of narrow passage. E: Detailed view of the column with stigma (s), pollinia (p) and anther cap (ac). Scale bar: 5 mm

The main goal of our study is to uncover the pollination strategy used by *Corybas* Remutaka, a taxonomically undescribed orchid endemic to NZ. Towards this end, we observed the orchid-pollinator interaction *in situ* and *ex situ*, studied the life cycle of the pollinator, and investigated the presence of olfactory and visual cues on the flowers, as well as micro-morphological features of the flower.

## MATERIAL AND METHODS

### The orchid

The identity of the orchid we studied may be verified by reference to the vouchers SP104171 and SP104172 lodged at WELT herbarium (Museum of New Zealand Te Papa Tongarewa, Wellington, NZ) and Genbank accessions KX499831 (nrITS), KX523933 (*trnL–trnF*), KX456127 (*psbJ–petA*) and KX499808 (nrITS), KX523931 (*trnL–trnF*), KX456129 (*psbJ–petA*).

### Pollination experiments, floral manipulation and pollinator observations

Pollination experiments were conducted during three flowering seasons between late winter and early spring (August–September). Plants with floral buds were covered using hydroponic plastic pots with the bottom wrapped in tulle so air circulation was allowed but floral visitors excluded. Once the flowers opened (ca. two weeks after exclusion), these were hand-pollinated and enclosures kept in place for another two weeks to prevent interference from potential floral visitors. Hand pollination treatments included cross-pollination (flowers were emasculated and then hand-pollinated with pollen from a plant 5–10 m away) and self-pollination (plants were hand-pollinated using their own pollen). We also tested for automatic self-pollination (plants were left intact and covered up until floral senescence), agamospermy (flowers were emasculated and covered up until floral senescence) and natural pollination (flowers were tagged but left uncovered).

To investigate whether the long filiform sepals and petals, a common feature of many NZ *Corybas,* are involved in the attraction of pollinators, we removed these from 20 flowers growing in the wild and later assessed natural fruit set. Sepals and petals were removed at the bud stage when they are tightly curled at each side of the bud.

Insect visitation was observed directly and recorded with a Sony Handycam (HDR-XR 160). The camera was left unattended recording floral visitors while pollination experiments were being conducted at different sites. Floral visitors were captured and later identified by Richard Toft, chief entomologist at Entecol Ltd. (www.entecol.co.nz). We conducted nocturnal observations, using a night vision camera, during the first flowering season but since no floral visitors were observed these were subsequently abandoned.

To investigate the diversity of fungus gnats (*Mycetophila*, Diptera) at the study site we set up a single Malaise trap (a tent-like trap) at about 5 m from where pollination experiments were set up. The trap was operated during the entire flowering season, i.e. mid-July to mid-October. The trap contents were recovered at five different times and examined under a dissecting microscope. Identification of *Mycetophila* was done following Tonnoir and Edwards (1927) and it was based on wing patterns and terminalia. All examined material was deposited at the entomological collection of the Museum of New Zealand Te Papa Tongarewa and available on-line via https://collections.tepapa.govt.nz/.

### Rearing studies and observation of the pollinators ex situ

Insect eggs were collected from the orchid labellum during early spring (late August – early September). These were placed in between the gills of white button mushrooms (*Agaricus campestris*) purchased from a local supermarket. The mushroom was kept in a glass jar, resting on a bed of sterilised forest leaf-litter mixed with fine gravel. The mouth of the jar was then covered with paper tissue and held in place by a rubber band. Jars were kept in a cold and humid environment and sprayed with water if the leaf litter appeared dry. Jars were checked daily to follow the development of the insects. Adult gnats were then used for the *ex situ* observations described below.

Adult insects were released inside a transparent Perspex acrylic box (500 mm^3^) where an orchid (leaf and flower only) was held in an Eppendorf tube with tap water. The box was kept in a cold and humid room and the video camera was left unattended filming for most of the day (0800 – 1800 h). The insects’ behaviour was observed from the time they emerged to their death.

### Floral micromorphology and olfactory and visual signals

To detect the presence and distribution of scent glands 10 fresh flowers and leaves were soaked in neutral red (1:10000) (Stern *et al*., 1986) for 2 h, and then washed in tap water and examined under the dissecting microscope. Scent glands should stain deep red (Stern *et al*., 1986). The micromorphology of the labellum, petals and sepals was also examined using scanning electron microscopy (SEM) and light microscopy (LM). For SEM observations, floral parts were preserved in ethanol (70 %). At the Manawatu Microscopy and Imaging Centre (Massey University, Palmerston North) samples were critical point dried, sputter-coated with gold and observed and photographed using a Cambridge 250 SEM.

Visual patterns were explored by photographing the flowers with a Nikon D70 DSLR camera, which has a sensor that is somewhat sensitive to UV light, and a EL-Nikkor 80 mm f5.6 enlarger lens, which admits UV wavelengths and focuses similarly in both visible and UV light (Schmidt, 2011). One set of close-up rings was used. First, the flower was photographed in normal light, then a Baader U Filter (UV-pass) was added and the exposure repeated. The UV image was illuminated by a Vivitar 285HV flash, modified by removal of its UV filter. The photographs were adjusted for brightness and the UV image desaturated to monochrome.

The false colour images were created using the software GIMP (https://www.gimp.org). First, the colour and UV photographs were cropped so they would overlap when combined. The colour photograph was decomposed into its red, green, and blue channels as separate layers, and the UV photograph superimposed as a fourth layer. Then the layers were recomposed, replacing red with the original green, green with the original blue, and blue with the original UV layer (Garnock-Jones, 2017). The resulting image approximates the light patterns seen by an insect, but in human-visible wavelengths, much as a piece of music transcribed into a lower key retains the relationships between the notes, but at a different pitch. However, the pixels of the two exposures have not always perfectly overlapped, leading to slight ghosting in some parts of the false colour images.

### Scent analyses

The orchid scent was investigated by solid-phase microextraction (SPME) and solvent extraction. The first procedure was used to detect volatile compounds (VOCs) emitted by the entire plant. Solvent extraction, on the other hand, allowed for detection of compounds present in specific parts of the plant (e.g. labellum, column and leaf). Unlike SPME, solvent extraction can detect large chemical compounds such as long hydrocarbon chains and many other non-volatile compounds.

SPME and solvent extraction analyses were conducted using a Shimadzu QP2010-Plus gas chromatograph mass spectrometer (GC-MS). The GC was fitted with a Restek RTX – 5SilMS column (30 m × 0.25 mm i.d. × 0.25 μm film thickness) and connected to an electron impact mass spectrometer, operating at 70 eV in positive ion mode, scanning from m/z 40–600 every 0.3 sec. The compounds were separated in the GC column with an initial oven temperature of 50 °C for 2 min, followed by a ramp at 10 °C / min to 300 °C, with a final hold for 5 min. Compounds were annotated by comparison of both retention time and mass spectra with those within the NIST 2008 and in-house libraries, and published spectra. Compounds present at similar abundance as in the controls were considered to be contamination and excluded from further analysis.

### SPME sampling

The entire plant (leaf and flower) was placed in a 20 mL glass SPME vial with septum and stored in a fridge at ca. 4° C for 12–18 hrs before measurement. This was done to concentrate any aroma emitted by the orchid. The vial was then incubated in a water bath at 20 °C for 20 min. This gentle heating increases the vapour pressure of the VOCs emitted by the plant and facilitates the absorption process of the volatiles by the SPME fibre. VOCs emitted by the plant were then collected using a SPME StableFlex^TM^ fibre kit (50 /30 μm DVB/Carboxen/PDMS coated fiber, needle size 24 ga). VOC extraction was performed by exposing the fibre inside the vial for 40 min while heating in the water bath. After adsorption the fibre was retracted into the needle, the syringe was removed from the vial and transferred to the GC-MS for sample injection. Volatiles adsorbed onto the fibre were desorbed within the splitless injection port of the gas chromatograph at an injector temperature of 270 °C using Helium as the carrier gas at a linear velocity of 40.1 cm / s (1.10 mL/min) at constant flow. The exposed fibre was always cleaned between samples for 2 min at 270 °C in the GC injection port. We used readings from empty clean vials and leaf only as controls.

### Solvent extraction

Different parts of the plant (entire plant, leaf, labellum and column) were immersed in 5 mL of the organic solvents Ethyl Acetate (EtOAc), Dichloromethane (DMC) and n-Hexane for 60 min at room temperature. After removing the plant material, the solvent was transferred into a 2 mL glass vial for GC-MS injection. Samples were introduced (0.5 μL) using a Shimadzu AOC-20i auto-sampler with a splitless injection mode at an injector temperature of 270 °C using Helium as the carrier gas at a linear velocity of 40.1 cm/s (1.10 mL / min) at constant flow.

## RESULTS

### Pollination treatments and floral manipulation

*Corybas* Remutaka did not form fruits by autogamous self-pollination or in the absence of pollen (Table 1). Fruit were only formed by flowers that were hand cross- or hand self-pollinated (90 % and 75 %, respectively). Natural fruit-set was low and only 12 % of the 79 unbagged flowers in the field formed fruit (Table 1). Flowers that had their sepals and petals removed also had low fruit-set; only one (5 %) of the 20 flowers formed fruit.

**Table 1:** Pollination experiments conducted *in situ* on plants of *Corybas* Remutaka across three flowering seasons (n= 165).

| Pollination treatment | Nos. flowers | Nos. fruit (%) |
| --- | --- | --- |
| Agamospermy | 11 | 0 |
| Automatic self-pollination | 21 | 0 |
| Hand self-pollination | 32 | 24 (75 %) |
| Hand cross-pollination | 22 | 20 (90 %) |
| Natural pollination | 79 | 12 (15 %) |

### Pollinator observations

The only floral visitor observed during the three seasons was the endemic fungus gnat *Mycetophila phyllura* (Mycetophilidae), despite 20 other species of *Mycetophila* that were present at the time of flowering **[Supplementary Table S1]**. Both male and female fungus gnats were observed on the flowers and captured carrying pollinia (Fig. 2). The behaviour these insects exhibited in the wild is described below and recorded here [**Supplementary Information - Video 1**]. Normally a male fungus gnat would land on the leaf or flower and either 1) sit still for minutes and then fly away, 2) explore the flower and then fly away, or 3) explore the flower (outside and inside) and position itself at the centre or side of the flower and start fanning its wings. This movement is accompanied by the regular up and down movement of the abdomen and opening of its claspers. If a female fungus gnat responds to this mating call, copulation briefly takes place (ca. 15 secs). The female will then either 1) fly away immediately or 2) explore the flower and probe the surface of the labellum with its ovipositor. If another male fungus gnat lands in or near the flower, the resident male displays a territorial behaviour and chases the recent arrival away.

**Figure 2:**
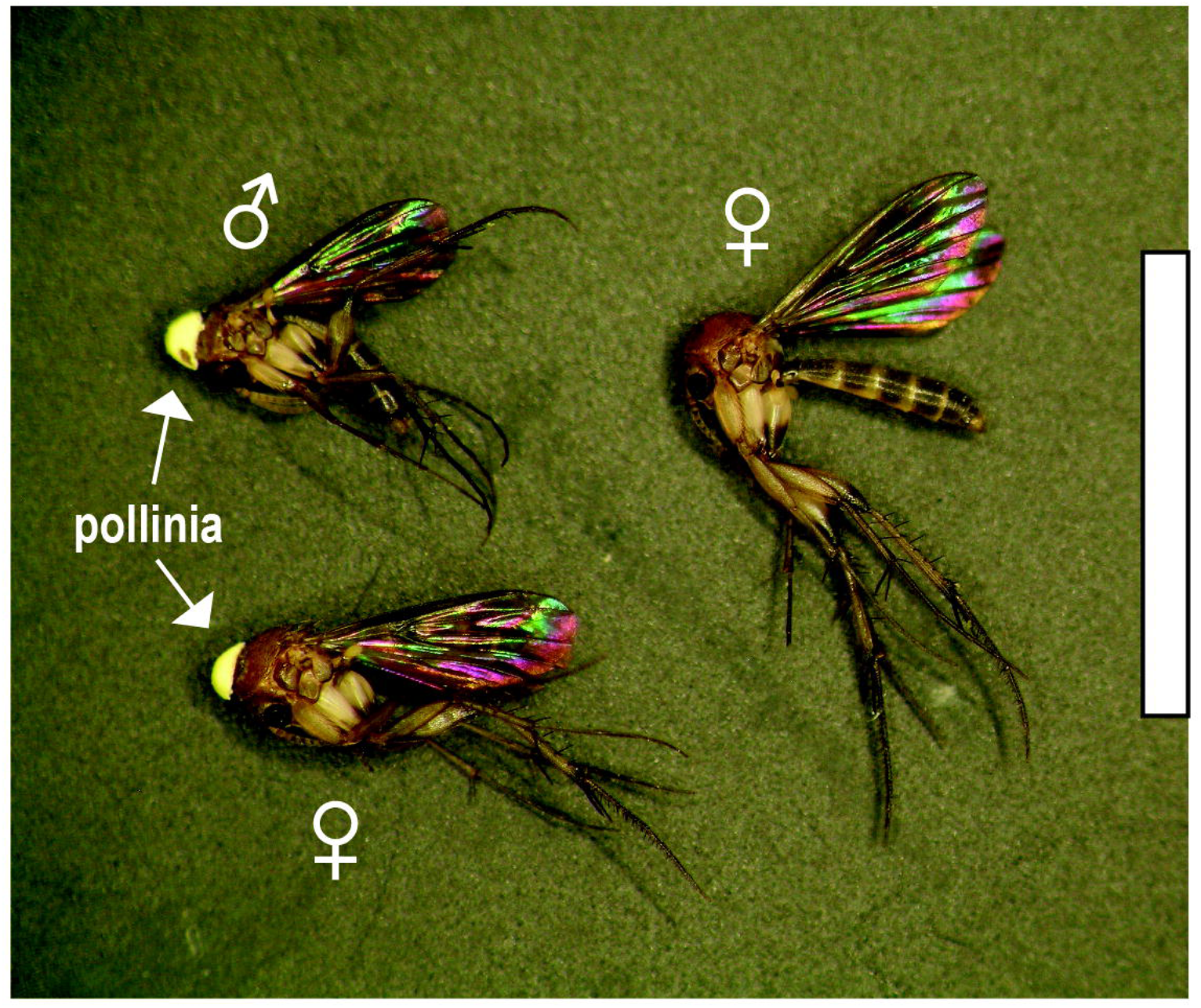
Male and female fungus gnats *Mycetophila phyllura* (Mycetophilidae) carrying pollinia (arrow) of *Corybas* Remutaka attached to their thorax. Scale bar: 5 mm

### Rearing studies and observation of the pollinators ex situ

After rearing these insects for two seasons, we can confirm the entire life cycle of *M. phyllura* takes about 30 days. After placing the eggs on the mushroom gills, it took about three days for the larva to hatch and about six days for it to leave the mushroom and form a cocoon among the mixture of leaf litter/pebbles at the base of the jar. After 10 days, the first adults emerged. We also noticed there was ca. three days delayed between adults emerging and becoming sexually active. During this time, insects remained mostly inactive, perching behind the orchid leaf or the box walls for hours. Courtship and mating behaviour was observed for ca. three days only, after this male and female gnats died.

Fungus gnats reared *ex situ* exhibited the same behaviour as those observed in the wild when presented with virgin flowers cultivated *ex situ*. Insects were mostly active in the mornings (from 0900 h until midday) and then again late afternoon (after 1600 – 1900 h). Fungus gnats gathered around the flowers, males displayed courtship behaviour, and copulation and oviposition were observed on the flower (Fig. 3A). Also, a female fungus gnat was observed crawling inside the flower, becoming stuck and later dying in there.

**Figure 3:**
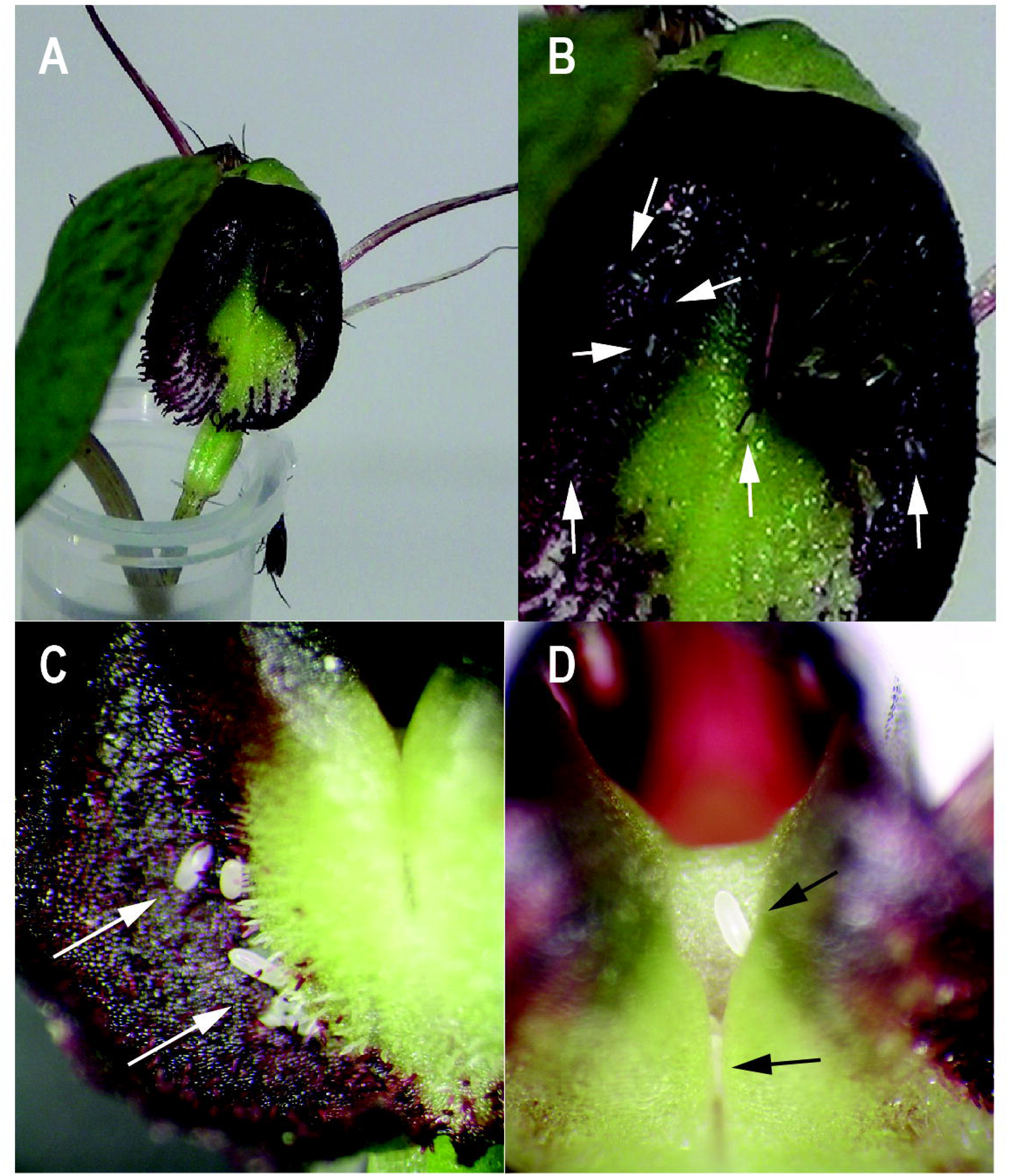
Fungus gnats (*Mycetophila phyllura*) gathering and mating on a flower of *Corybas* Remutaka *ex situ* (A, B). Insects were reared under controlled conditions from eggs founds on the labellum of plants growing in the wild (C, D). Arrows point to eggs laid on the labellum.

### Scent glands and UV patterns

Both the flower and part of the leaf showed a positive reaction to neutral red stain. In the flower, a positive reaction was observed along the upper surface of the dorsal sepal near the apex, the distal section of lateral petals and sepals and the hairs on the labellum lamina (Fig. 4). Inside the flower, the anther cap and the stigma reacted positively to the stain; the stigmatic surface however is normally absorbent so this result was ignored. Only the leaf apicule stained deep red.

**Figure 4:**
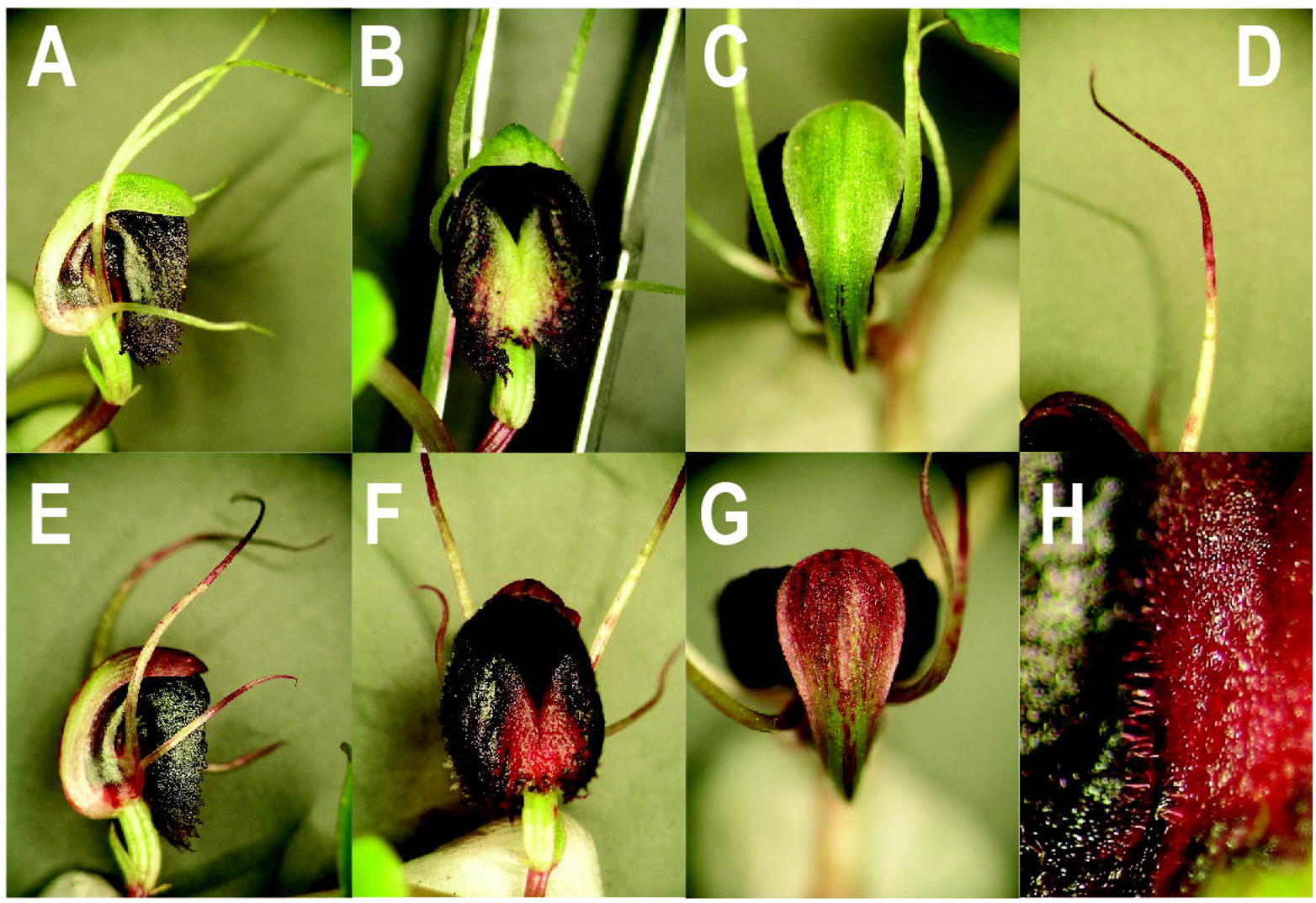
Flower of *Corybas* Remutaka before (A-C) and after (D-H) soaking in a bath of neutral red for 2 hr. The presence of putative scent glands (osmophores) is indicated by the uptake of the red colour.

We did not observe specific UV patterns, but UV reflection was stronger at the distal end of the ovary and base of the dorsal sepal (near the auricles), the middle section of the petals, the external surface of the labellum right under the dorsal sepal and internal surface of the labellum (Fig. 5).

**Figure 5:**
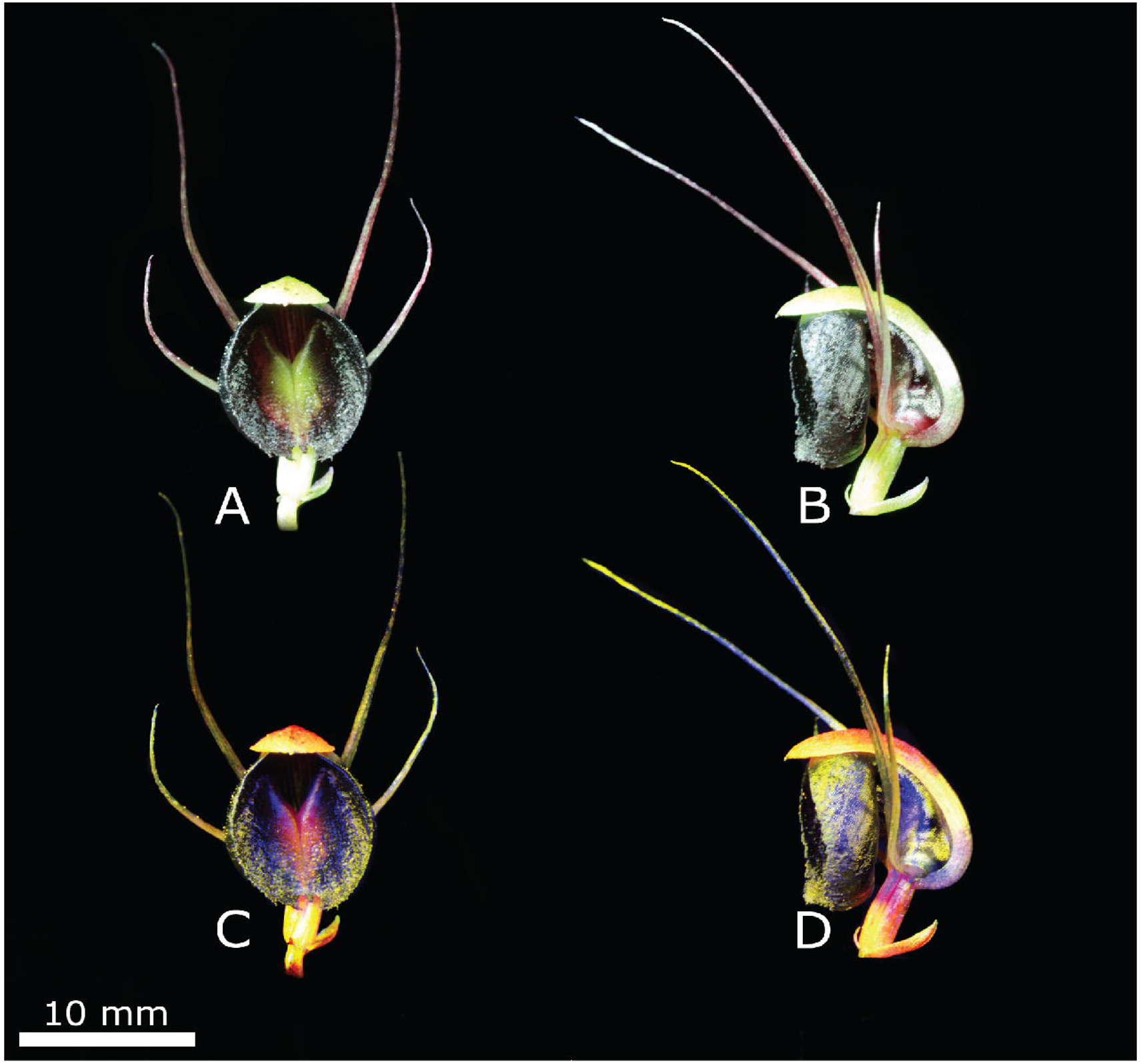
Frontal and side view of flowers of *Corybas* Remukata (A, B) and their false colour images (C, D). Areas of most intense uv reflection appear blue.

### SEM observations

Microscopic observations revealed that the surfaces of both lateral sepals and petals are smooth, formed by elongated rectangular cells and with no evidence of secretory glands (Fig. 6). The upper surface of the dorsal sepal, on the contrary, is formed by cells of irregular shape with a number of round to ovoid-shaped cells, solitary or in clusters, scattered along the vascular bundles which run longitudinally from the base of the petal to the apex, where they were more abundant (Fig. 6C). Of all floral structures examined, the labellum’s upper surface was the most complex (Fig. 6D) and, based on its micromorphology, at least four areas can be distinguished. These are described below from the lateral margins to the center of the labellum, followed by the proximal section (towards the base of the column) and upper section (above the column). A number of unicellular trichomes, solitary or in clusters of 2–4 (Fig. 6E), were observed mostly along the main vascular bundles of the labellum. These are accompanied by round to ovoid cells similar to those observed on the apex of the dorsal sepal. Trichomes are more abundant towards the center of the labellum than the margin, and they become strongly appressed, pointing towards the center of the flower (i.e. towards the column) (Fig. 6F). Along the central furrow of the labellum, and walls facing the column, the surface is scale-like, with their walls slightly raised. A number of fine ridges were observed on the surface of each of these scale-like papillae. The labellum surface then transitions from scale-like papillae to elongated and rectangular cells (Fig. 6G). These occur towards the lateral margin of the labellum, which joined tightly at the back of the column and above the column. Despite the smooth appearance of these cells, their surface is ornate with ridges that run, mostly in parallel, towards the centre of the flower (i.e. the column) (Fig. 6H). Finally, the top surface of the anther cap is densely papillose and the surface of each papilla decorated with fine ridges.

**Figure 6:**
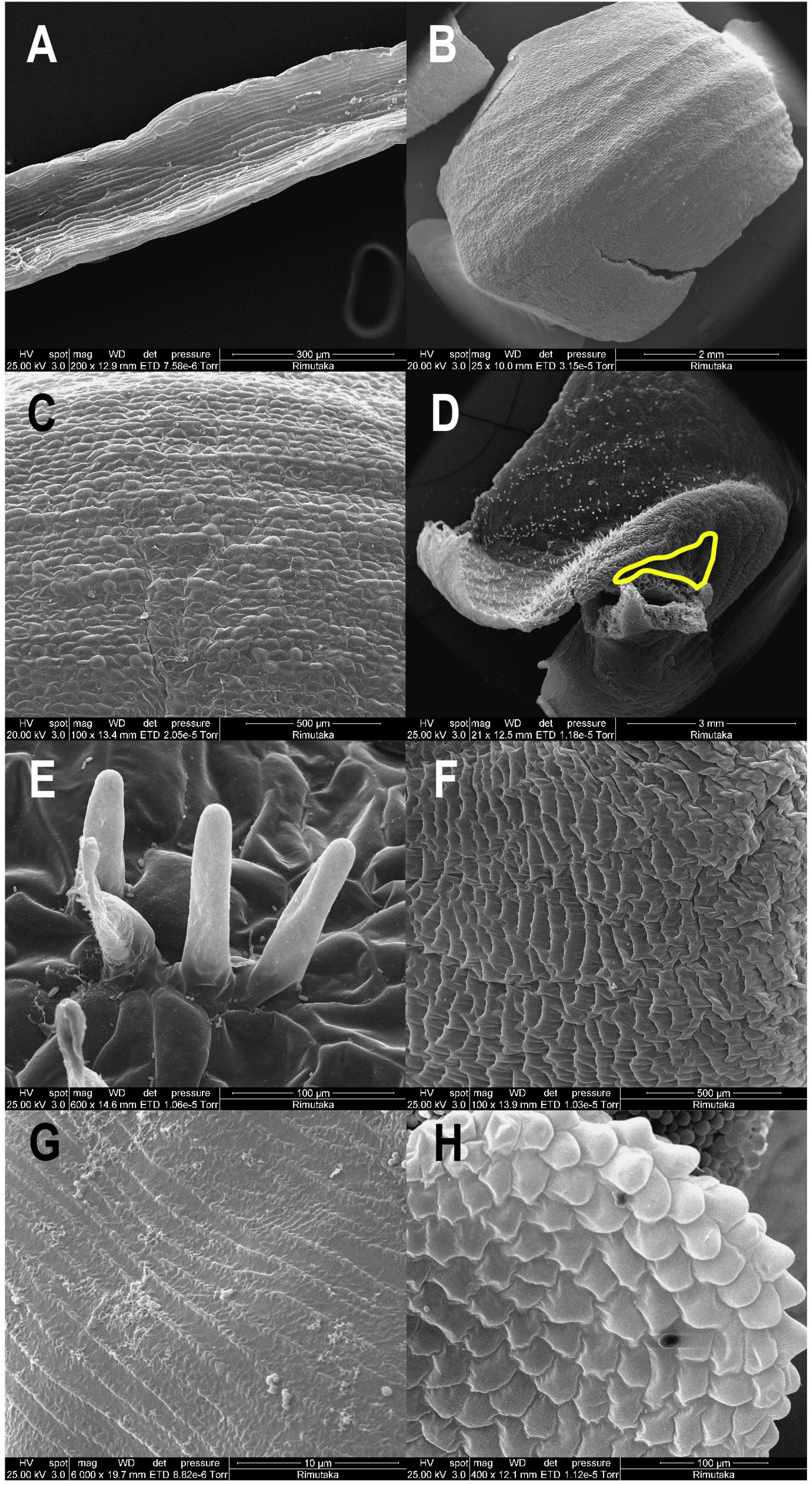
Photographs of the floral parts of *Corybas* Remutaka as seen under the scanning electron microscope. A: Sepal. B: Dorsal sepal. C: Details view of dorsal sepal. D: Labellum upper surface. Yellow line delimits the depression in the labellum where the female taps with its ovipositor and eggs are sometimes found. E: Close up view to trichomes found on the labellum. F: Scale-like papillae found in the central part of the labellum. G: Details of the labellum surface right above the column. H: Close up view to the papillae on the externals surface of the anther cap.

### SPME sampling

Altogether, the GC-MS analysis detected 21 compounds from the four orchid samples studied by SPME (Table 2). These peaks represent eight chemical classes with alkane, fatty acid derivatives and sesquiterpenoids being the most abundant (Table 2). The number of peaks detected per sample ranged from nine to 11 and only three compounds were detected across all four samples; these were cyclooctanyl acetate, copaene and 9,12,15-octadecatrienal (Table 2). The next most common compounds were hexadecane, cycloisosativine and an undetermined alkane found in only three of the four samples (Table 2). Overall, the concentration of these and remaining compounds varied considerably between samples but the sesquiterpene cycloisosativene, the fatty acid derivative cyclooctanyl acetate and the monoterpene α-pinene had the highest relative abundance across samples (Table 2).

**Table 2:** Compound annotations detected by solid-phase micro-extraction (SPME) and their concentration (area / %) across four samples (leaf and flower) of *Corybas* Remutaka. RT: Retention time. —: Not detected.

| Chemical group | RT (mins) | Annotation | Sample 1 | Sample 2 | Sample 3 | Sample 4 |
| --- | --- | --- | --- | --- | --- | --- |
| Amine | 2.22 | 2-Pentanamine | 0.61 | — | — | — |
| Other cyclic lipids | 3.54 | Tetrahydrofuran | — | — | 0.55 | — |
| Monoterpene (Bicyclic) | 5.26 | α-Pinene | 3.18 | — | — | — |
| Benzenoid | 6.82 | 1-Methoxy-4-methyl-benzene | — | 1.39 | — | — |
| Fatty acid derivative (Acetate ester of fatty alcohol) | 7.99 | Cyclooctanyl acetate | 1.81 | 2.12 | 2.47 | 3.58 |
| Benzenoid | 10.44 | 2,4-Dimethoxytoluene | 2.42 | 2.00 | — | — |
| Fatty acid derivative | 11.60 | Methyl 13,16-octadecadienoate |  | 0.21 | — | — |
| Sesquiterpene (tricyclic) | 12.14 | Cycloisosativene | 5.56 | — | 5.34 | 2.84 |
| Sesquiterpene (tricyclic) | 12.21 | Copaene | 0.79 | 0.34 | 0.16 | 0.18 |
| Alkane | 12.46 | Dodecane | 0.12 | 0.22 | — | — |
| Other cyclic lipids | 12.80 | Spiro[2.4]heptane, 1,2,4,5-tetramethyl-6-methylene- | — | — | — | 0.93 |
| Sesquiterpene (Bicyclic) | 12.95 | trans-α-Bergamotene | — | — | 2.51 | 0.19 |
| Menthane monoterpene | 13.05 | Isoterpinolene | — | — | — | 0.23 |
| Monoterpene (monocyclic) | 13.24 | o-Mentha-1(7),8-dien-3-ol or 3,3-Dimethyl-hepta-4,5-dien-2-ol | — | — | 0.48 | 0.64 |
| Sesquiterpene (acyclic) | 13.61 | β-Farnesene | — | — | 0.10 | — |
| Alkane (C16) | 13.75 | Hexadecane | 0.31 | 0.34 | — | 0.64 |
| Fatty acid derivative (Acetate ester of fatty alcohol) | 15.78 | cis-7-Dodecen-1-yl acetate | — | — | — | 0.64 |
| Fatty alcohol (unsaturated) | 17.97 | 9,12,15-Octadecatrienal | 0.29 | 0.28 | 0.50 | 0.64 |
| Sesquiterpene (cyclic) | 18.72 | 1R,3Z,9s-4,11,11-Trimethyl-8-methylenebicyclo[7.2.0]undec-3-ene | — | 0.27 | — | — |
| Alkane (C24) | 20.19 | Undetermined | — | 0.22 | 0.30 | 0.64 |

### Solvent extraction from entire plant and individual parts

Overall, 19 peaks were detected by solvent extraction of plants (flower + leaf) *in situ* and *ex situ* (Table 3). Annotation of these compounds indicate they are mostly alkanes and fatty aldehydes. Independently of the solvent used, the chemical profiles of samples extracted *in situ* and *ex situ* were very similar. A few exceptions were the presence of tetradecane detected in samples extracted in the field, and azulene detected only in two of the three samples extracted in the laboratory (Table 3). The compound with the highest relative abundance in the field extractions was tetradecane (Table 3). Meanwhile, heneicosane and hexatriacontane were the most abundant compounds found in samples extracted in the lab. These were also the only two compounds detected in all labellum extracts (Table 3). Many of the compounds detected in the column extracts were the same as those recovered from *in situ* and *ex situ* extractions with the exception of squalene, which was detected when using the solvents EtOAc and Hexane (Table 3). From all compounds detected in the column, the alkene 9-hexacosene had the highest relative concentration along with 2,5,8,11-tetraoxadodecane which was detected only when EtOAc was used as solvent (Table 3). Finally, leaf extracts were dominated by tetradecanal and hexatriacontane.

**Table 3:**
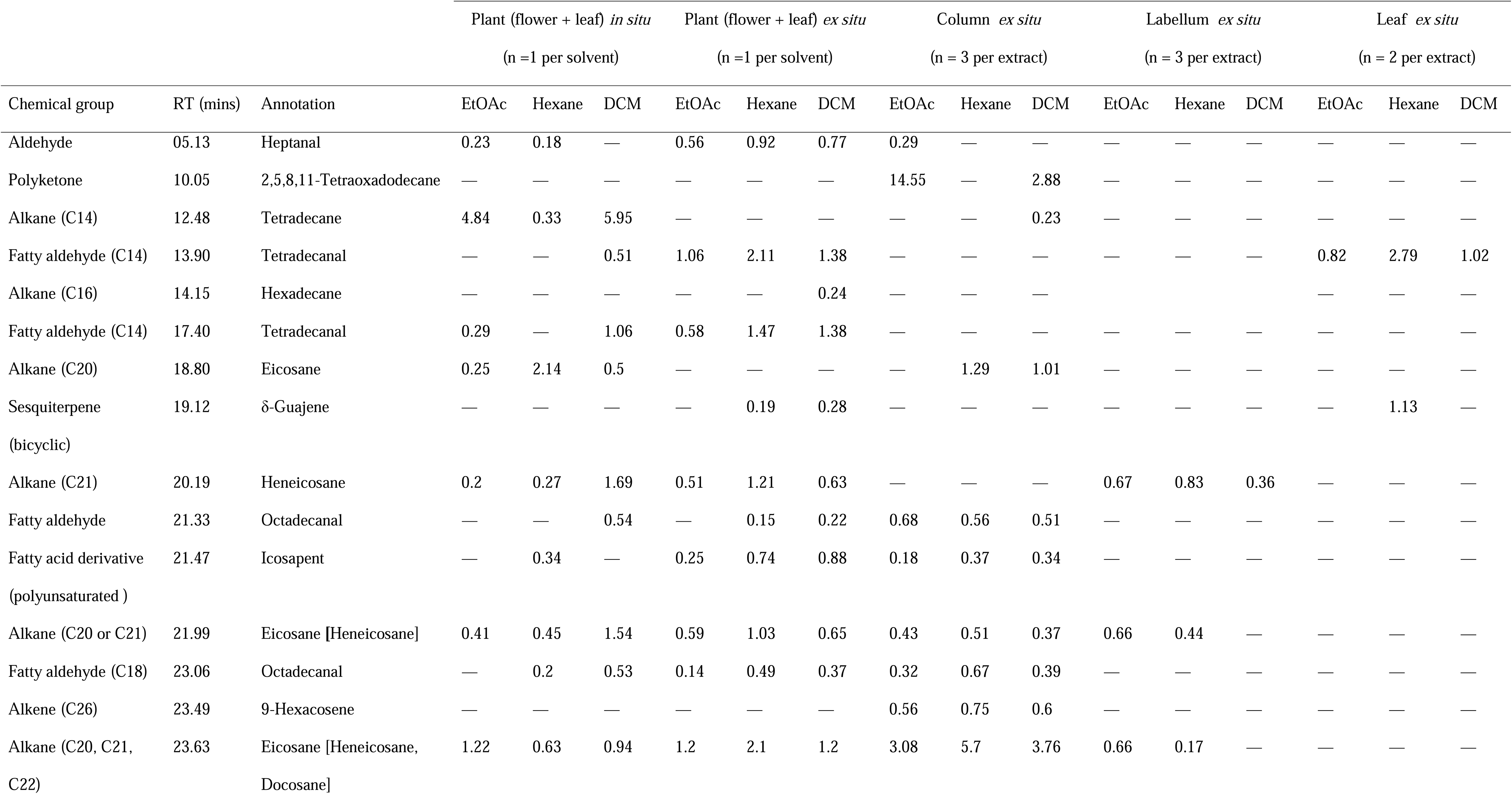
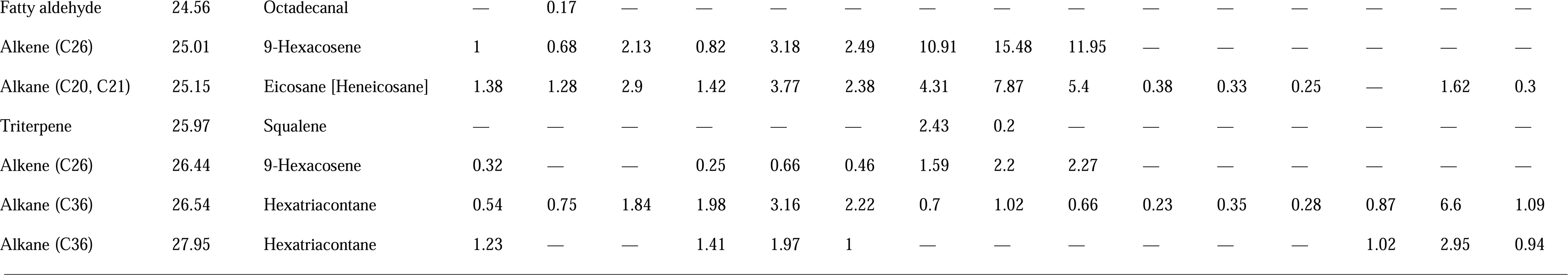
Compound annotations detected in *Corybas* Remutaka plants by solvent extraction conducted in the field (*in situ*) and in the laboratory (*ex situ*) and their concentration (area / %). Compounds indicated in brackets are second and third most likely identifications. RT: Retention time. —: Not detected. Number of carbons are indicated in parenthesis.

### Mushroom solvent extraction

Altogether 76 peaks were detected from the mushroom where fungus gnats were observed either mating or laying eggs **[Supplementary information - Video 2]**. The best-represented chemical groups were alkanes, fatty alcohols, and fatty acid derivatives (Table 4). Compounds in common with orchid samples were eicosane, 9-hexacosene, squalene, hexadecane and hexatriacontane. From all the compounds detected in this mushroom, eicosane and 17-pentatriacontene had the highest relative abundance (Table 4).

**Table 4:**
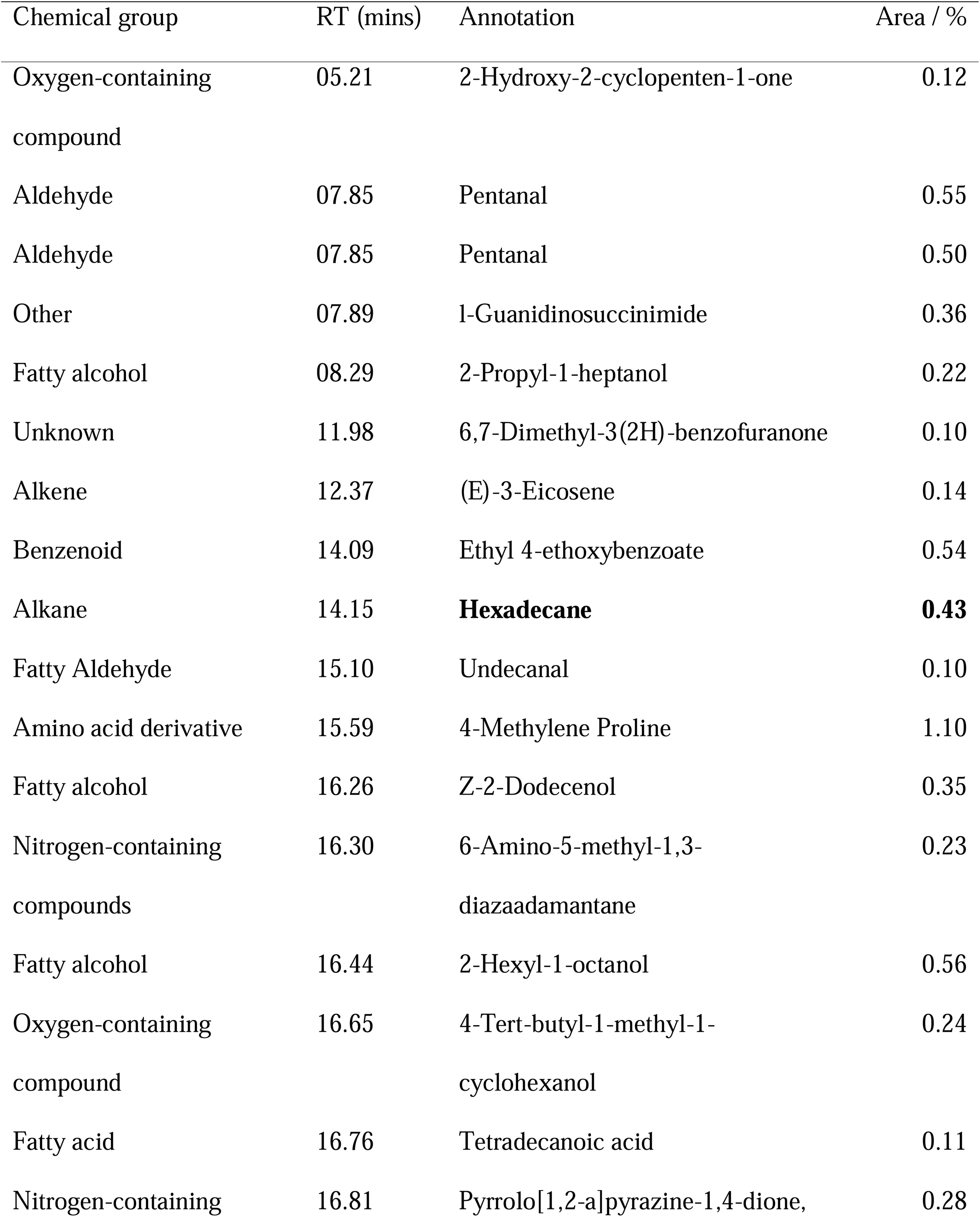
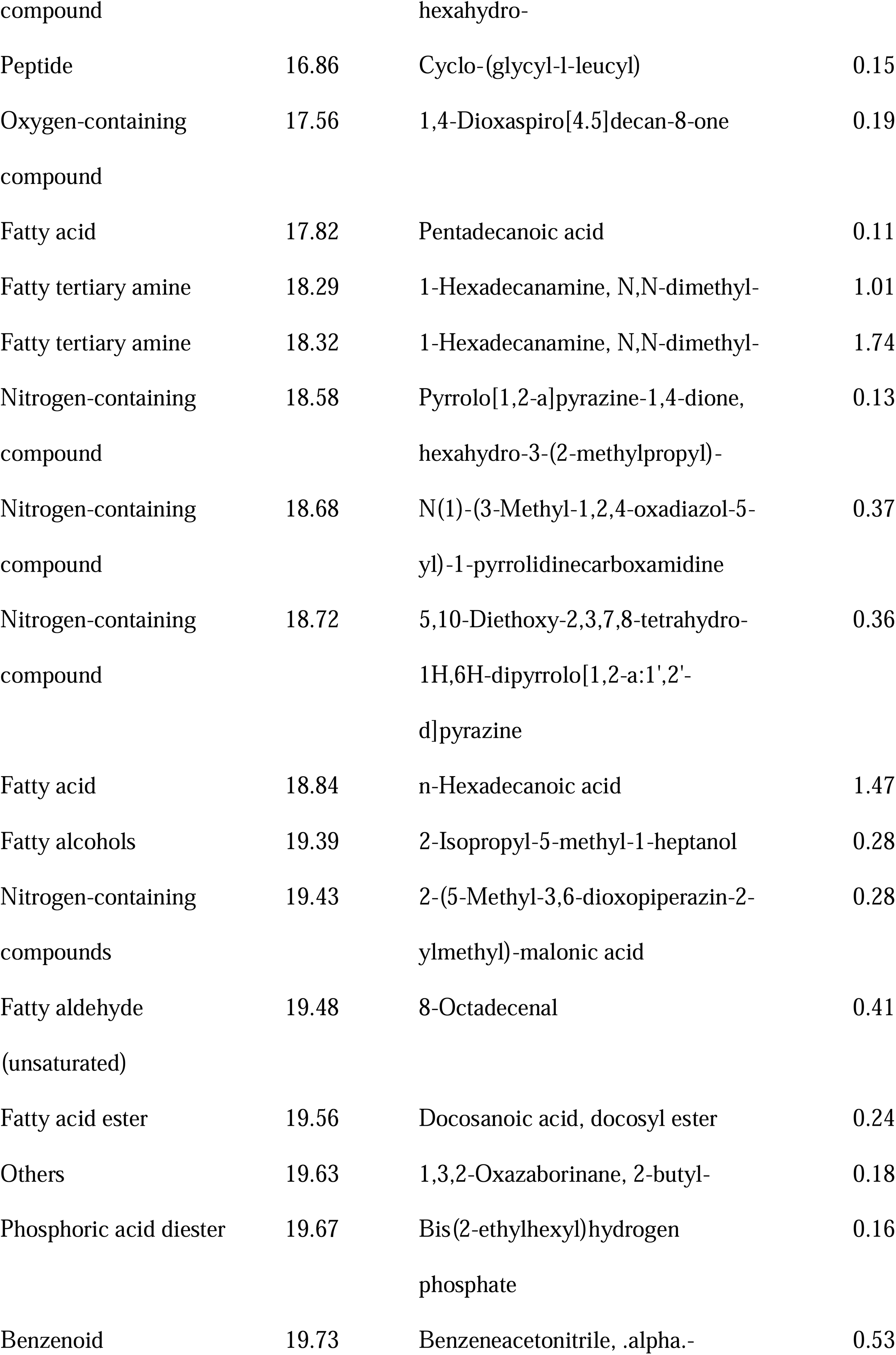
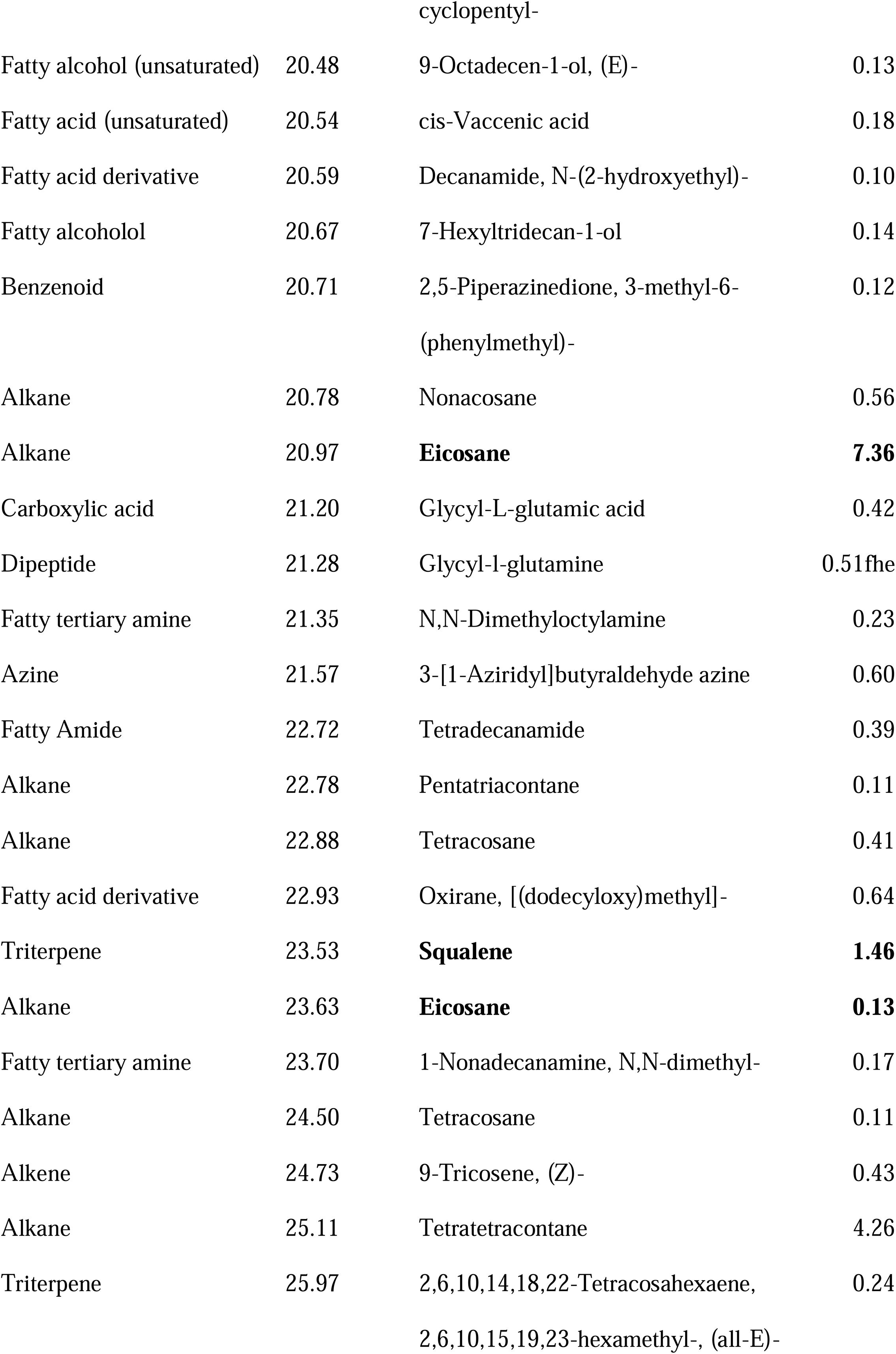
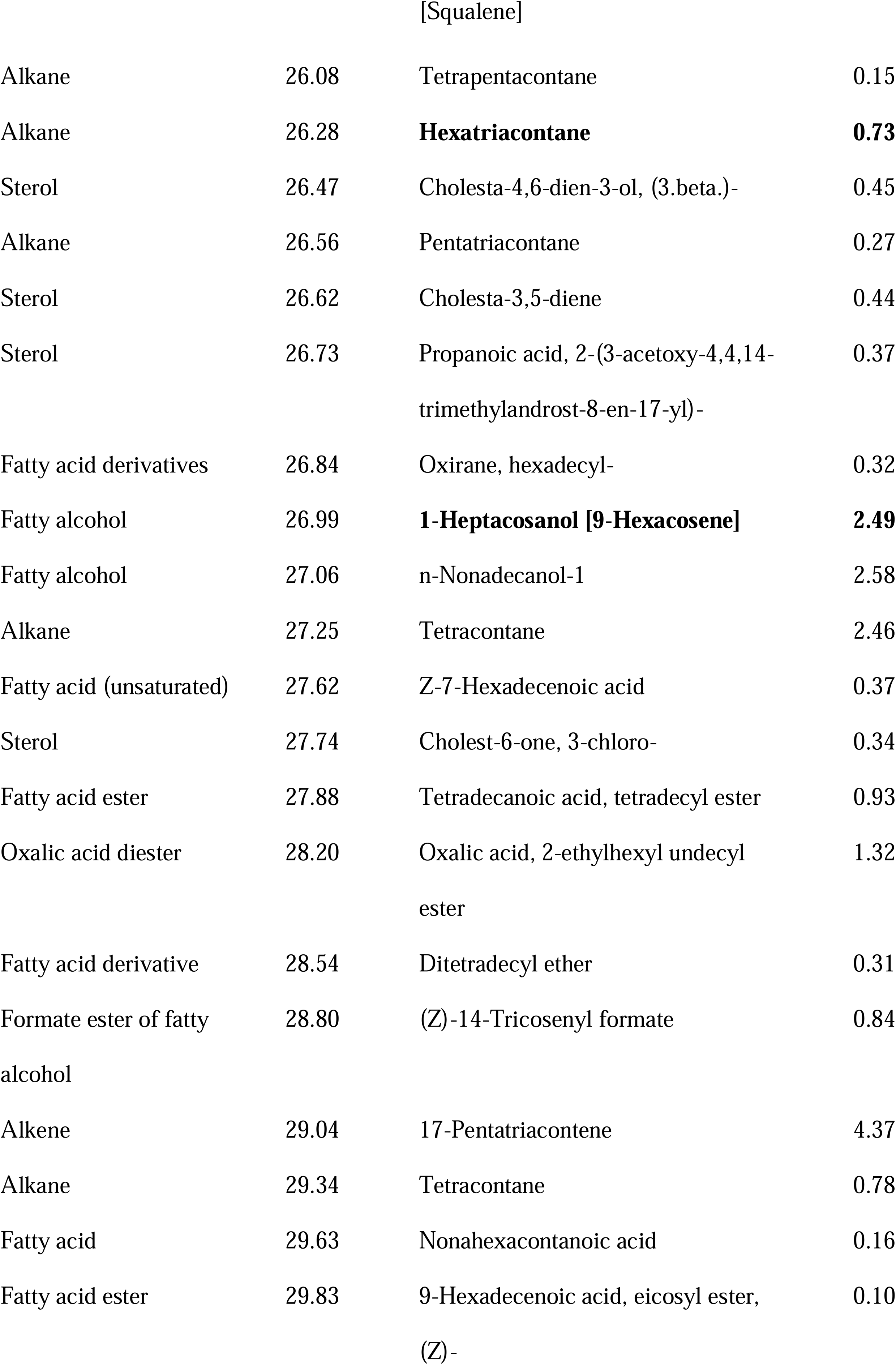
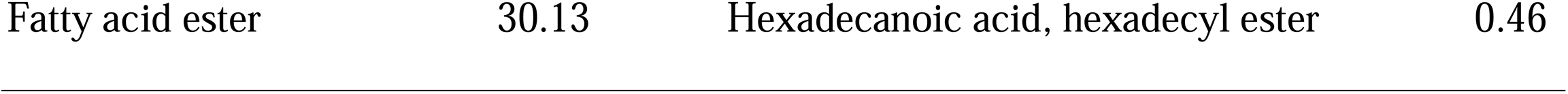
Compound annotations detected by Hexane solvent extraction in the laboratory of the fruiting body of a single individual of an unidentified mushroom found at one of the study sites and visited by *Mycetophila phyllura*. Compounds also detected in *Corybas* Remutaka are indicated in bold.

## DISCUSSION

### Breeding system, pollinators and morphological adaptations

Our pollination experiments have demonstrated *Corybas* Remutaka is a nectarless orchid incapable of self-pollination that depends on a specific insect, the fungus gnat *Mycetophila phyllura* (Diptera), for the fertilisation of its flowers. In our study, only those flowers that were hand self- or hand cross-pollinated formed capsules, 75 % and 90 % fruit-set, respectively (Table 1). These values are in strong contrast with the 15 % fruit-set produced by natural pollination suggesting pollen limitation (i.e. low visitation rate) rather than resources is limiting fruit-set. Low pollination rates are common in non-rewarding orchids. For instance, Neiland and Wilcock (1998) reported the global mean fruit-set for non-autogamous nectarless orchids in temperate areas is 28.5%. Values lower than the global mean, such as 1.2–4%, have been reported in other NZ orchids in the genus *Corybas* (Darwin, 1890 but Kelly *et al*., 2013) and *Pterostylis* (Lehnebach *et al*., 2005; Bodley *et al*., 2016 but Thalwitzer *et al*., 2018). Although the absence of a reward is the most obvious reason for low visitation rate, local overabundance of the rewardless species, so pollinators learn to avoid it (Moret *et al*., 1998), competition for the same pollinator by rewarding species, and rarity or absence of the pollinator (Wilcok and Neiland, 2002) may also reduce pollination rates considerably. All three reasons may apply to *Corybas* Remutaka, but pollinator abundance/rarity is the most likely explanation. Our survey of the fungus gnat diversity at the study site recorded 21 species of *Mycetophila* and although we captured more than 1700 individuals **[Supplementary Data, Table S1]**, only one specimen belonged to *M. phyllura*. It is unlikely this result is an artefact of our trapping method as Malaise traps are commonly used to collect fungus gnats in NZ and overseas (e.g. Harris *et al*., 2004; Jakovlev *et al*., 2014) but a reflection of the fungus gnat biology. For instance, their distribution across landscapes is generally patchy due to their reduced dispersal ability (Jakovlev, 2012) and their abundance irregular as they rely on the presence of mushrooms, which are seasonal, to complete their life cycle as their larvae feed on them only (Jakovlev, 2012). Furthermore, earlier holdings of this species in the entomological collection at the Museum of New Zealand Te Papa Tongarewa (https://collections.tepapa.govt.nz/taxon/82906) suggests this species is widespread but locally uncommon.

Observations *in situ* and *ex situ* have confirmed that both sexes of *M. phyllura* visit the flowers of *Corybas* Remutaka, transport pollen and therefore act as pollinators (Fig. 2) **[Supplementary information - Video 3]**. These findings contradict earlier claims that only female fungus gnats visit and pollinate *Corybas* (van der Pijl and Dodson, 1966; Dafni and Bernhard 1990). The behaviour displayed by these insects on the orchid **[Supplementary Information, Video 1]** followed this sequence: 1) male gnat lands on the leaf or center of the labellum; 2) male gnat inspects the flower and chamber formed by the folding of the labellum around the column; 3) male gnat starts wing fanning with up and down movement of the abdomen; 4) female gnat lands on the leaf or flower and slowly approaches the male; 5) insects mate on the flower or leaf; 6) after mating, the female either explores the flower and probes the labellum surface and a depression at the top of the labellum central groove (see Fig. 6D) with its ovipositor or flies away right after mating. The male stays on the plant and resumes with the behaviour mentioned in step 3. Male or female gnats can remove/deposit pollinia during steps 2 and 6. Fungus gnats reared under controlled conditions displayed this same behaviour when presented with the orchid plant. Female fungus gnats even laid eggs on the labellum (Fig. 3B) but a few days after hatching the larvae died, as there is no food for them in the flower. Only larvae moved to mushroom fragments reached adulthood. Besides our study, we have found only one other example of a non-rewarding plant species pollinated by fungus gnats which oviposit on the flowers; i.e. *Asarum tamaense* (syn. *Heterotropa tamaensis*) (Aristolochiaceae) (Sugawara, 1988; Kakishima and Okuyama, 2020).

To the best of our knowledge, our study is the first to describe the courtship and mating behaviours of NZ fungus gnats, their life cycle, and behaviour while visiting *Corybas* flowers. This is not surprising as this pollination event is subject to multiple variables and therefore not commonly observed. For instance, observations have to coincide with the time *Mycetophila* are more active, mornings and dusk (Jakovlev, 2012), time when these short-lived insects are sexually active (restricted to only ca. three days) and suitable weather conditions which are uncommon in late winter-early spring. Furthermore, the orchid should occur within the home range of the pollinator. Presently, it is unknown how restricted and specific the distribution and habitat preferences of *M. phyllura* are, but it is likely *Corybas* Remutaka may have extended its occurrence beyond the home range of its pollinator. At our study site this orchid is extremely abundant forming large patches of 20 or more plants growing in very close proximity, most likely propagated via clonal reproduction (van Etten *et al*., 2018).

Some of the behaviours we have described above have also been observed, and stunningly illustrated, on a range of Australian *Corybas* also visited by fungus gnats (Kuiter and Findlater-Smith, 2017). The main difference to our observations is that Australian fungus gnats were reported as feeding on a “secretion” produced by the labellum surface and a “concentrated viscous liquid” produced at the base of the column (Kuiter and Findlater-Smith, 2017). No signs of such secretions nor feeding behaviour were observed in *Corybas* Remutaka nor in any other NZ *Corybas*. Overall, our observations provide no support for the presence of a reward in NZ *Corybas* as has been suggested in Australian species (Jones, 1970; Kuiter and Findlater-Smith, 2017).

Another event that we observed under controlled conditions was fungus gnats becoming trapped headfirst inside the flower in a space formed by labellum walls (Fig. 1D). This passage is formed by the lateral margins of the labellum tightly joined. Dead fungus gnats have been found imprisoned inside the flowers of many other NZ *Corybas* (Upson, 2014) and even a new species of fungus gnat was described in NZ from a specimen imprisoned inside the flower of *Corybas oblongus* [syn. *Corysanthes oblonga*] (Miller, 1918). These insects appear to become glued by their thorax to the stigma or viscidial disc. In their struggle for freedom the gnats may pollinate the flower: self-pollinate if the pollinia has not been removed and the flower has not been already fertilised, or cross-pollinating it if the insect is carrying pollinia. The auricles, the two lateral openings at the base of the flower (Fig. 1B), may lure the insect, positioned headfirst, deeper into this tight space as they may appear as potential exit holes. Backing out from this tight passage is unlikely due to modifications of the upper surface of the labellum. The entrance to this passage and the lateral walls have scale-like papillae all leaning inwards; i.e. towards the center of the flower (Fig. 1D and 6F). Meanwhile, cell walls at the roof of the passage have ridges running lengthwise with arrow-like ornamentation also pointing towards the center of the flower (Fig. 6G). Similar surface modifications, including exit holes, have been described from trap-flowers (or kettle flowers) which secure their fertilisation by trapping their pollinators (*e.g*. Vogel and Martens, 2000; Bröderbauer *et al*., 2012) and the pitchers of carnivorous plants (e.g. Adams and Smith, 1977; Schwalliger *et al*., 2020).

### Olfactory and visual signals

For a long time, pollination of *Corybas* has been associated with brood-site deception and it has been suggested flowers smell and look like mushrooms (e.g. van der Pijl and Dodson, 1966; Dafni and Bernhard, 1990; Jersakova *et al*., 2006; Kelly *et al*., 2013; Kelly and Gasket, 2014). Annotation of the compounds detected in *Corybas* Remutaka shows a prevalence of alkane, sesquiterpene and fatty aldehydes (Tables 2, 3). None of the volatiles we detected here were those typically reported from brood-site deceptive flowers mimicking carrion (sulphur compounds) (Urru, 2011), faecal and urine (phenol and indole derivates, acids) (Urru, 2011), fermentation odours (small aliphatic alcohols and esters) (Martos *et al*., 2015; Stökl *et al*., 2010) or mushrooms (alcohols and ketones) (Policha, 2014; Policha *et al*., 2016). More specifically, compounds such as 3-octanone, 3-octanol and 1-octen-3-ol, which are commonly found in mushrooms and other mushroom-mimics such as the orchid *Dracula lafleurii* (Policha *et al.,* 2016), were missing. Typical fungal compounds were also missing from the putative brood-site deceptive *Corybas chessemanii* (Kelly and Gaskett, 2014). To some extent, this is not surprising because unlike *Dracula lafleurii* neither of these two *Corybas* smell like mushroom to the human nose. Interestingly, these compounds were also missing from the mushroom where *M. phyllura* were observed mating (Table 4) which may suggest the occurrence of 3-octanone, 3-octanol and 1-octen-3-ol is not widespread across fungi.

We have found, however, a few compounds shared by the orchid and the mushroom that may be involved in the attraction of fungus gnats. One of them is a 36-carbon compound annotated as hexatriacontane. This compound was detected by solvent extraction of the whole plant (Table 3) and extracts from individual parts such as leaf, labellum and column (Table 3). Hexatriacontane has been found in the surface wax of plant leaves (Li and Ishiwaka, 2006; Rahman *et al*., 2019; Mobarak *et al*., 2020), flowers (Mukherjee *et al*., 2013; Jakubska-Busse *et al*., 2014), mushrooms (Mohamed and Farghaly, 2014; Milovanovic *et al*., 2020) and the cuticular hydrocarbons of insects (Qadir *et al*., 2021). A recent study has shown long-chain alkanes (i.e. C_15_-C_36_) like hexatriacontane play an important role as short-range attractants for insects to find their host and as ovipositional stimulant (Mobarak *et al*., 2020). Other long-chain alkanes found in the orchid and the mushroom were hexadecane (C_16_) and eicosane (C_20_), and the triterpene squalene. Again, these are compounds that serve as attractants or sexual pheromones that may trigger oviposition and probing behaviour in insects **[Supplementary Table S2]**. Squalene has been found in the NZ bat pollinated holoparasitic plant *Dactylanthus taylorii* (Ecroyd *et al*., 1995). Further to these four compounds, we detected a number of semiochemicals (i.e. chemicals involved in the biological communication between insects) which are known to trigger a range of reproductive behaviours in Diptera, Hymenoptera and Lepidoptera, ranging from aggregation, lek-like, pre-mating and mating behaviour to oviposition [**Supplementary Table 2**]. Some examples are copaene which triggers lek-like behaviour and guides insects to *rendezvous* sites (Nishida *et al*., 2000) and tetradecanal, a putative component of the sex pheromone of tachinid flies recently found in the sexually deceptive orchid *Telipogon peruvianus* (Martel *et al*., 2019).

Our findings suggest some or a blend of these compounds may be responsible for the attraction of *M. phyllura* and its behaviour, but further research is needed to confirm this. Future studies should use synthetic standards to validate compound identification, and behavioural and electrophysiological tests are needed to determine compounds’ biological activity. Studies should also explore scent composition and emission at different times of the day and flower lifespan, as these may vary considerably (e.g. Chapurlat *et al*., 2018). Daily variation in floral scent emission may also explain the absence of typical fungal compounds in *Corybas* as both studies, i.e. Kelly and Gasket (2014) and this study sampled scent at a single moment in time. The same applies to scent analyses of mushrooms, because the onset of sporulation or the death of a fruiting body may strongly affect the quality and quantity of the odour bouquet (Fäldt *et al*., 1999). In addition, sampling should ideally happen *in situ* as some compounds may degrade and not be detected after the plants are moved into laboratory conditions (e.g. greater concentration of tetradecane detected in field extractions than those in the lab, Table 3).

Similarly, further characterisation of scent glands in *Corybas* Remutaka is required. Scent emission is likely to occur at the apex of the dorsal sepal, the upper surface of the labellum, anther cap (Fig. 4) and, interestingly, also the leaf apicule (not shown). Although these structures had a positive reaction to neutral red and their micro-morphology resembles to that observed in the osmophores of other orchids (e.g. Kowalkowska *et al*., 2015), further histological characterisation are required to confirm their function, especially so for the leaf apicule, as there were no obvious secretory structures observed there under LM and SEM.

Besides olfactory signals, there are number of visual cues linked to fungal mimicry. These include floral traits such as a brown or green perianth with white stripes or a perianth with flagelliform appendages (Johnson and Schiestel, 2016). Both traits occur in the flowers of *Corybas* Remutaka (Fig. 1) and many of the remaining NZ species (Lehnebach *et al*., 2016). However, whether the dull colour of these orchids mimics the colour of co-occurring mushrooms is debatable as analyses of spectral reflectance data for the putative mushroom-mimic *C*. *cheesemanii* showed no evidence for colour mimicry but strong UV reflectance (Kelly and Gasket, 2014). We also detected UV reflectance in *Corybas* Remutaka, but not at the top of the dorsal sepal as in *C. cheseemanii*. UV reflectance was observed in the middle section of petals, along the center of the labellum (seen as two parallel bands running longitudinally) and near the auricles (*i.e.* exit holes) (Figure 5D). Our findings provide further support to Kelly and Gasket’s (2014) idea that *Corybas* may attract fungus gnats by exploiting their sensory bias for UV. In *Corybas* Remutaka, UV reflectance may work as a close-range cue for fungus gnats after landing on the flower and serve as a guide towards the stigma and exit holes at the base of the column. However, we believe that UV reflectance may not be as relevant as the olfactory signals discussed above as the removal of the long filiform UV-reflecting sepals did not prevent visitation or fertilisation (1 out 20 flowers formed fruit).

### Pollination strategy

Based on the evidence we have gathered in this study we believe that *Corybas* Remutaka rely on a mixed strategy that combines elements of *rendezvous* site attraction and brood-site deception. Thus, males and female fungus gnats meet and mate on the flowers (*rendezvous* site attraction), and gravid gnats oviposit on the flowers (brood-site deception). Similar to brood-site deception, *rendezvous* site attraction is a deceptive mechanism where a plant secure pollination of its flowers by exploiting the sexual response of its pollinator (Dafni, 1984). In this system, both sexes of a pollinator meet in the flower and subsequently mate (Faegri and van der Pijl, 1979).

*Rendezvous* site attraction has been reported in European and African orchids pollinated by bees and beetles, respectively (see summary by Jersakova *et al*., 2006) and *Dracula* orchids from Ecuador (Endara *et al*., 2010). The labellum of *Dracula* orchids look and smell like mushroom, and attract a number of drosophilid mycophilous flies, which use the flowers as a *rendezvous* site and shelter during the rainy season (Endara *et al*., 2010). These flies court and mate on the flowers but unlike what we have observed in *Corybas* Remutaka, neither probing behaviour by female insects nor eggs have been found on the flowers (Endara *et al*., 2010). Furthermore, and unlike *Corybas* Remutaka, *Dracula* orchids offer a sort of reward, a liquid film produced by the labellum on the first day the flower opens (Endara *et al*., 2010).

Pollination by fungus gnats (e.g. Mycetophilidae and Sciaridae) occurs in several plant families, including orchids (see citations in Reiter *et al*., 2019). Their attraction to orchid flowers involves rewarding and deceptive strategies such as the presence of nectar (e.g. Jones, 1974; Ackerman and Mesler, 1979), sexual deception (Blanco and Barbosa, 2005; Phillips *et al*., 2014; Reiter *et al*., 2019) and a combination of food reward and brood-site deception (Duque-Buitrago *et al*., 2014). The latter strategy is somewhat similar to that described in *Dracula* orchids (see Endara *et al*., 2010) and although Duque-Buitrago *et al*. (2014) observed mating and oviposition behaviour, no eggs were found in the flower as we did in *Corybas* Remutaka (Fig. 3C, D). Another interesting difference between our observations and those by Duque-Buitrago *et al*. (2014) is that in *Corybas* Remutaka, male fungus gnats were always the first to land on the orchid and then, after performing a series of “calls” (i.e. wing fanning and up-and-down moving of the abdomen), female gnat(s) arrived. This suggests males are the primary target of this pollination strategy and females are later attracted by the calls and/or pheromones male fungus gnats make and release from the orchid.

This apparently obligate relationship between an orchid and its only pollinator has likely consequences for both. First, a requirement for effective mimicry by the orchid is likely to be under strong selection, because orchids that do not deceive a gnat will not reproduce. Secondly, a requirement for a gnat’s ability to distinguish a good host (mushroom) from a bad one (orchid) is also potentially under strong selection, because deceived fungus gnats will not reproduce (unless they can mate again on a mushroom; even then their fecundity is likely reduced). But the orchid is also dependent in the long term on maintenance of a fungus gnat population. Attraction of only male fungus gnats to the flowers might indicate a role for sexual selection: perhaps male fungus gnats are less selective than females are the about quality of mating sites.

## Conclusion

Our study shows that *Corybas* Remutaka is a self-compatible, nectarless, pollinator-dependent orchid that relies on a mix of volatiles and micro-morphological features to attract a specific pollinator, the fungus gnat *Mycetophylla phyllura*. The orchid is a true specialist and exploits the reproductive instinct of male and female fungus gnats to secure pollination. This deceptive pollination strategy combines features of *rendezvous* site pollination and brood-site deception. Our study is the first to thoroughly document the pollination biology of *Corybas*, the life cycle of a native fungus gnat, and unequivocally demonstrate the occurrence of this specialised and deceptive pollination system in the NZ flora. Since specialised pollination systems like this are linked to increased species diversification (Schiestl and Schluter, 2009; Sinn *et al*., 2015) it would not be surprising that future studies uncover this unusual partnership has promoted reproductive isolation within *Corybas* and, ultimately, driven its diversification in NZ.

## ACKNOWLEDGEMENTS

We thank George and Keena Gibbs for allowing us to conduct this study at their property and the Greater Wellington City Council for research permits to work at local regional parks and forests. This work was supported by the Marsden Fund [MNZ1001 to C.A.L].

## SUPLEMENTARY DATA

**Table S1**: Species of *Mycetophila* (Diptera) collected at the study site using a Malaise trap.

**Table S2**: Compounds detected in *Corybas* Remutaka and unidentified mushroom and their biological activity as reported in The Pherobase (https://www.pherobase.com/) and literature.

**Video 1**: Fungus gnats (*Mycetophila phyllura*) landing and calling from flowers of *Corybas* Remutaka.

**Video 2**: Fungus gnats (*Mycetophila phyllura*) mating on unidentified mushroom found near *Corybas* Remutaka flowers.

**Video 3**: Male fungus gnat (*Mycetophila phyllura*) carrying pollinia and calling from flowers of *Corybas* Remutaka.

## AUTHOR CONTRIBUTIONS

C.A.L conducted the pollination experiments, floral characterisation, fungus gnats rearing experiments and *ex situ* observations of pollinators. A.J.Z filmed pollinator’s behaviour in the wild and lead scent analyses with contribution of R.K. J.K. helped with insect collection and identification and PG-J conducted the UV analyses of the flowers. C.A.L lead the writing with contributions from the other authors.

